# A simulation-based approach to improve decoded neurofeedback performance

**DOI:** 10.1101/450403

**Authors:** Ethan Oblak, James Sulzer, Jarrod Lewis-Peacock

## Abstract

The neural correlates of specific brain functions such as visual orientation tuning and individual finger movements can be revealed using multivoxel pattern analysis (MVPA) of fMRI data. Neurofeedback based on these distributed patterns of brain activity presents a unique ability for precise neuromodulation. Recent applications of this technique, known as decoded neurofeedback, have manipulated fear conditioning, visual perception, confidence judgements and facial preference. However, there has yet to be an empirical justification of the timing and data processing parameters of these experiments. Suboptimal parameter settings could impact the efficacy of neurofeedback learning and contribute to the ‘non-responder’ effect. The goal of this study was to investigate how design parameters of decoded neurofeedback experiments affect decoding accuracy and neurofeedback performance. Subjects participated in three fMRI sessions: two ‘finger localizer’ sessions to identify the fMRI patterns associated with each of the four fingers of the right hand, and one ‘finger finding’ neurofeedback session to assess neurofeedback performance. Using only the localizer data, we show that real-time decoding can be degraded by poor experiment timing or ROI selection. To set key parameters for the neurofeedback session, we used offline simulations of decoded neurofeedback using data from the localizer sessions to predict neurofeedback performance. We show that these predictions align with real neurofeedback performance at the group level and can also explain individual differences in neurofeedback success. Overall, this work demonstrates the usefulness of offline simulation to improve the success of real-time decoded neurofeedback experiments.

## 1 Introduction

Multi-voxel pattern analysis (MVPA) extracts information about a person’s cognitive state by analyzing spatially distributed patterns of functional MRI activity (Lewis-Peacock and Norman, 2014; Haxby et al., 2014). This approach has become ubiquitous in cognitive neuroscience since the seminal work of Haxby et al. (2001) identified distributed and overlapping representations of visual object categories in temporal cortex. MVPA is especially useful for isolating fine-grained relationships between brain activity and behavior, such as orientation tuning (Kamitani and Tong, 2005; Haynes and Rees, 2005) and complex motor programs (Wiestler and Diedrichsen, 2013; Kornysheva and Diedrichsen, 2014) which are inaccessible to other human neuroimaging analysis methods. However, despite the increased signal-detection sensitivity of MVPA, conventional neuroimaging research is limited in its ability to draw causal inferences about brain-behavior relationships.

Investigation into causal mechanisms of MVPA representations of neural activity requires this activity to be modulated. However, techniques such as TMS (Walsh and Cowey, 2000) and tDCS (Brunoni et al., 2012) are incapable of modulating fine-grained patterns of neural activity. Operant conditioning of neural activity, known as neurofeedback, uniquely enables self-modulation of a target neural circuit through feedback, most often presented visually (Sulzer et al., 2013; Sitaram et al., 2017). Early work in fMRI neurofeedback mirrored contemporary univariate techniques in offline fMRI analysis (Ruiz et al., 2014). In recent years, MVPA-based neurofeedback techniques have taken hold (LaConte et al., 2007). For instance, a seminal work by Shibata et al. (2011) used neurofeedback based on decoded activity from early visual cortex, a process dubbed ‘decoded neurofeedback’ or ‘DecNef’. The researchers were able to show that individuals could learn to self-modulate a targeted pattern of brain activity related to a given orientation of a visual grating without stimulus presentation. Intriguingly this was associated with heightened perceptual acuity specific to the underlying stimulus. Thus, used in this manner, decoded neurofeedback is a powerful and unique tool in neuroscience that can manipulate neural activity patterns to reveal causal relationships with behavior. This technique has been used in several applications beyond low-level visual perception, including fear conditioning (Koizumi et al., 2017), confidence judgements (Cortese et al., 2016), and facial preference (Shibata et al., 2016).

It is well-known that a large proportion (up to 30%) of willing participants are unable to self-regulate their brain activity through neurofeedback training (Allison and Neuper, 2010; Hammer et al., 2012). The causes of this are not well understood, and the ‘non-responder’ problem remains a key challenge for neurofeedback research and clinical translation. Our previous work showed how different factors can affect decoded neurofeedback performance using a novel simulation paradigm (Oblak et al., 2017). Using feedback based on simulated brain activity in visual cortex, participants performed simple ‘cognitive strategies’ by choosing how to rotate an oriented grating clockwise or counterclockwise until a hidden target orientation was found. This simulation paradigm enabled the manipulation of ‘neurofeedback’ provided to the participant to reflect the signal quality and timing of realistic neural activity, and of unrealistic neural activity that would be impossible to present in the scanner by altering or removing the hemodynamic delay. Therefore we could link explicit strategy choices, and thus neurofeedback performance, with the characteristics of the feedback signal received. The approach produced insights into how fMRI neurofeedback presentation can enhance or inhibit learning; for example, intermittent feedback is better than continuously presented feedback when participants have a poor understanding of the hemodynamic properties of the brain signal. However, these simulations did not account for a key element in neurofeedback performance as it relates to decoded neurofeedback: the accuracy of decoding the desired fMRI activity patterns in real-time.

Decoding accuracy can vary widely between experiments and conditions. Standard processing techniques, such as normalization, detrending and averaging over time will all affect decoding accuracy (Hanke et al., 2009). However, to date there has been no systematic approach to investigating the effects of these parameters on neurofeedback performance. Numerous decoded neurofeedback studies lack an empirical justification for parameter selection (Watanabe et al., 2017), leaving the possibility of suboptimal neurofeedback training, which may contribute to the non-responder problem. The goal of the present study was to examine how real-time fMRI decoding accuracy is affected by these parameters, and likewise, how decoder accuracy contributes to neurofeedback performance.

Being able to address this question in a systematic manner requires explicit knowledge of neurofeedback strategies being used by participants. However, cognitive strategies used for neurofeedback, which commonly take the form of mental imagery (deCharms et al., 2005), can be difficult to identify and quantify. Here, we simplified this challenge by focusing on a restricted set of explicit strategies: individual finger movements, which are supported by neural correlates in primary sensorimotor cortex (M1/S1). There is ample literature on fMRI neurofeedback of mean regional activity in M1/S1 (Yoo and Jolesz, 2002; Weiskopf et al., 2004; Chiew et al., 2012; Friesen et al., 2017), and evidence that univariate M1/S1 self-modulation is associated with improvements in fine-motor control. For instance, Bray et al. (2007) found improved reaction time following M1/S1 neurofeedback training, and Blefari et al. (2015) observed a positive correlation between precision grip motor skill and neurofeedback performance. There are, however, no published attempts at decoded neurofeedback for M1/S1. Thus, by using a well-defined neural circuit (M1/S1) and measurable strategies (finger pressing), we sought to gain insight more generally into how people learn to use decoded neurofeedback, and under what conditions such learning is best facilitated.

Here, we designed a ‘finger localizer’ experiment based on previous finger individuation decoding experiments (e.g. Diedrichsen et al. (2012); Ejaz et al. (2015)) that could be used to train fMRI pattern classifiers for a neurofeedback experiment (Fig 1). We had participants (N=6) complete two of these localizer sessions, on separate days, which allowed us to investigate multiple parameters that impact real-time decoding performance in M1/S1. We first simulated participant neurofeedback performance using feedback yoked to individual localizer trials. This fully automated simulation helped us examine estimated neurofeedback performance in different conditions such as changes in region-of-interest (ROI) and different feedback success thresholds. Then, we recruited a new set of human participants (N=10) to perform this target-finding experiment with the same yoked neurofeedback and compared performance to participants in a real neurofeedback session. The combined simulation and experimentation provide a unique method for investigating fundamental questions about neurofeedback. The results may help to mitigate the non-responder problem, making decoded neurofeedback a more robust procedure.

**Figure 1:**
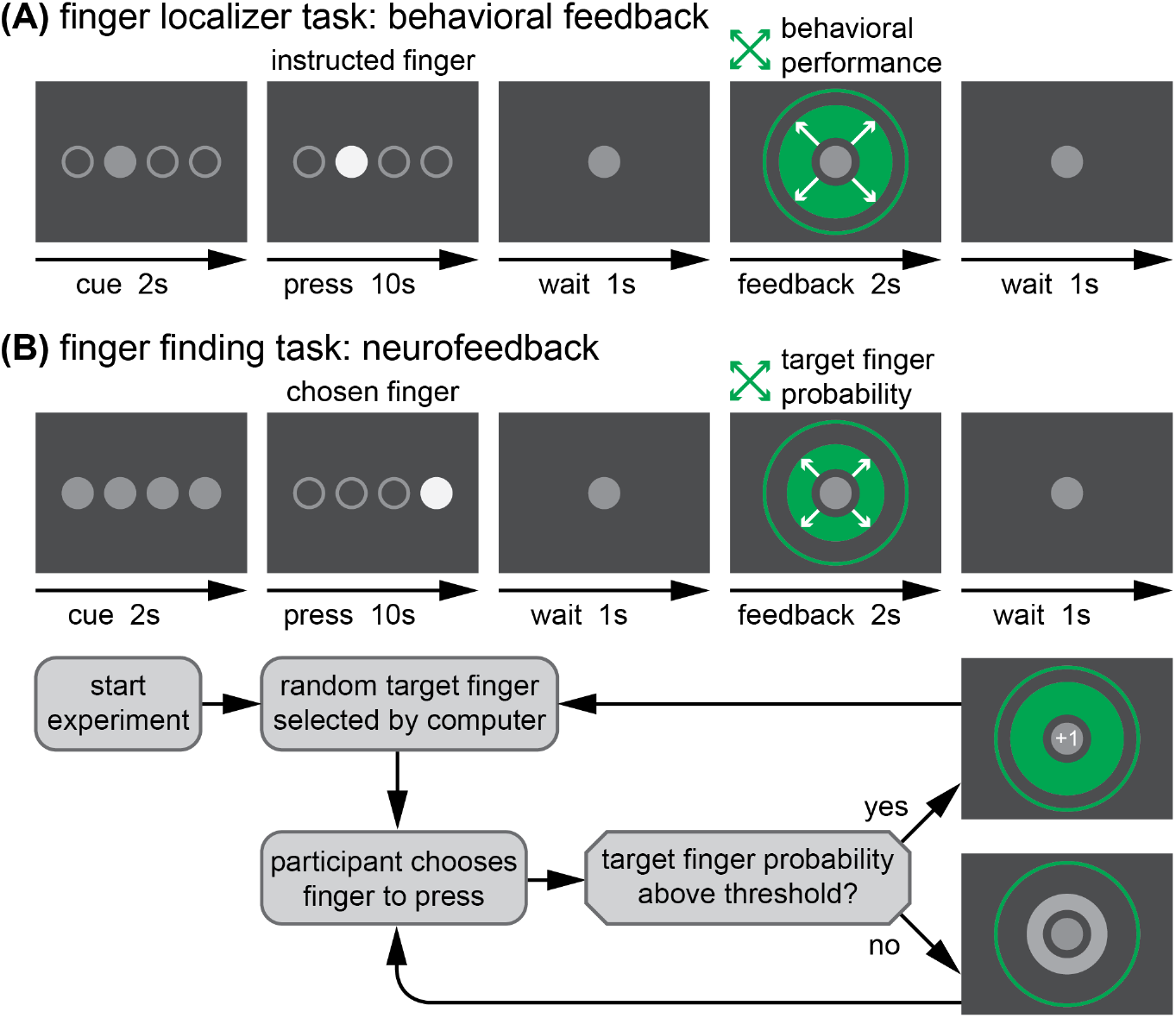
Experimental design. In both localizer **(A)** and neurofeedback **(B)** trials, a cue precedes a 10 sec period of finger pressing at 1 Hz, followed by feedback. The localizer feedback reflects behavioral performance for repeated presses of a finger chosen by the experimenter. The neurofeedback reflects the real-time fMRI decoder output for the target finger based on presses of a finger chosen by the participant. Below, the decision process for advancing trials in the neurofeedback session is presented. The target finger remains the same from trial to trial until a predetermined success threshold is reached, at which point a new random target finger is selected.

## 2 Materials and methods

### 2.1 Participants

Six healthy participants (2 female, average age 26.4 years, SD=2.4) were recruited from the University of Texas at Austin community in accordance with the University of Texas Institutional Review Board. All subjects completed two localizer sessions, but one subject was unable to participate in the neurofeedback session due to a hand injury that occurred after the second localizer session. An additional ten healthy participants (3 female, average age 24.3, SD=3.6) were recruited for a simulated neurofeedback experiment using the brain data from the localizer sessions of the original group of participants. Written informed consent was obtained from all participants.

### 2.2 General procedure

The experiment consisted of three fMRI sessions separated by at least 24 hours: two localizer sessions followed by one neurofeedback session. The localizer and neurofeedback sessions were similar in structure, as shown in Fig 1. The primary purpose of two localizer sessions was to identify the fMRI activity patterns in sensorimotor cortex corresponding to pressing each of the four fingers of the right hand (index, middle, ring, little). However, the secondary purpose of the localizer was to imitate the timing and processing limitations of a real-time fMRI neurofeedback session. Therefore, the duration of trials and runs were identical in both types of sessions, with both consisting of 8 fMRI runs. These finger-specific activity patterns identified in the localizer were used as targets in the neurofeedback session.

Each run began with a 40 sec baseline period, in which only a grey fixation circle was visible (diameter: 1.5° of visual angle). During the final 3 sec of the baseline period, the fixation circle flashed white at 1Hz to indicate the beginning of trials. Each run consisted of 20 trials of 16 sec each: 2 sec to cue the target finger, 10 sec of finger presses, 1 sec of rest, 2 sec of feedback, and 1 sec of rest before the next trial. On each trial, four circles appeared in the center of the screen (corresponding to the four fingers; 1.5° each, spanning 10° horizontally total) and were used to coordinate finger presses. In the cue period of the localizer task, only one circle turned grey indicating that this was the finger to be pressed on the trial. In the neurofeedback task, all circles turned grey indicating that the participant should choose one finger to press for the duration of that trial. Then, finger presses were cued at a rate of 1Hz for 10 sec with a filled white circle corresponding to the cued (localizer) or selected (neurofeedback) finger.

To ensure consistent behavior from participants, we encouraged rhythmic presses. Participants received positive visual feedback if presses occurred within a specific response window (200-500 ms) of each 1-sec epoch during the 10-sec pressing period. This response window was dynamically adjusted during the localizer sessions (see Section 2.3 for details), but remained constant for the neurofeedback session. The cued (or chosen) finger was filled in with a color corresponding to a participant’s performance on that press, beginning when the finger was pressed and ending at 800 ms into the 1-sec press epoch. Presses that occurred inside the response window filled the pressed finger’s circle green, and presses that were either too fast or too slow filled the circle yellow. If the incorrect (or unchosen) finger was pressed, the correct finger’s circle filled red. After a brief 1-sec wait period, trial-ending visual feedback appeared as a centrally presented green circle (2-10°) that expanded or contracted based on performance during the preceding 10-sec pressing period. In the localizer session, feedback was based on the rhythmicity of presses on that trial (i.e., the proportion of presses made within the desired response window). In the neurofeedback session, feedback was based on the correspondence between the pattern of fMRI activity for the target finger learned by the classifier during the localizer sessions, and the actual real-time pattern of fMRI activity evoked by the chosen finger on that trial. On the first trial, the feedback circle expanded smoothly from the origin to a diameter corresponding to the feedback value. At each subsequent trial, the feedback circle expanded or contracted over 500ms from the previous diameter to the updated diameter to ensure a smooth visual display. When the feedback on a neurofeedback trial exceeded the ‘target found’ success threshold, the starting point for feedback on the next trial was reset to zero and a new target was selected (see Section 2.4 and Fig 1B).

### 2.3 Localizer sessions (Days 1 and 2)

As stated earlier, the purpose of localizer sessions was to identify the fMRI activity patterns in sensorimotor cortex corresponding to pressing each of the four fingers of the right hand (index, middle, ring, little). The general task procedures for the localizer task are described above. All participants completed two localizer sessions separated by no more than 7 days (5+/-2 days, mean+/-s.d.). In each session, the 8 fMRI runs consisted of 20 total trials, with 5 trials for each of the 4 fingers. The order of trials was pseudorandomized to ensure an approximately equal number of each finger transitions (including pressing the same finger two trials in a row). Both localizer sessions were identical other than the order of button presses. As described above, we encouraged rhythmic presses by providing visual feedback (color-filled circles) for each finger press based on whether it was made within a desired response window (200-500 ms). In the localizer task, one ‘point’ was awarded for each correct response made within this window. Points were tallied at the end of each trial and mapped onto the green feedback circle, with 0 points corresponding to the minimum circle diameter and 10 points corresponding to the maximum diameter. The total score was also tallied and presented to participants at the end of each run. To control for task difficulty, an adaptive staircase procedure was used to incrementally adjust the rewarded response-time window based on performance after each trial. A threshold of 70% correct responses (7 points) was selected for staircasing. If this threshold was exceeded (8 or more points), then the upper limit of the time window decreased by 20 ms (i.e., to 480ms), making the task slightly harder. If performance was below this threshold (6 or fewer points), then the upper limit of the window increased by 20 ms (i.e., to 520ms), making the task slightly easier. If performance matched this threshold, no changes were made to the response window.

### 2.4 Neurofeedback session (Day 3)

The purpose of the neurofeedback session was to investigate whether human participants could efficiently and accurately interpret fMRI decoder outputs related to pressing the four fingers of the right hand on a trial-by-trial basis. As such, the participant’s goal was to respond to the decoded neurofeedback by finding and then pressing with the finger associated with the targeted brain pattern (i.e. ‘target finger’). Participants had 160 trials (20 trials per run, 8 runs total) to find as many target fingers as possible. A series of target fingers was pseudo-randomly generated for the experiment. A full set of 160 targets was generated in the unlikely case that a target was found each trial. The order of targets was determined by a concatenation of 20 lists that each contained a randomly shuffled arrangement of 8 finger targets (2 for each finger). This allowed for the target finger to occasionally repeat and to prevent prediction of the target finger (e.g. by process of elimination over a series of targets). Participants chose one finger to press each trial. After 10 presses of the same finger, the feedback circle appeared, as in the localizer sessions, except that here the size of the circle corresponded to the fMRI decoder output for the target finger in sensorimotor cortex (M1+S1, see Section 2.7). The decoder outputs (from 0 to 1, for each finger) indicated the likelihood estimates from the fMRI pattern classifier that the fingers were pressed. The minimum diameter of the feedback circle corresponded to 0% probability of the target finger, and the maximum diameter corresponded to 100% probability (Fig 1B). If the participant had been pressing the target finger during the trial, its output value should be relatively high. Note that the target output could also be spuriously high when a different finger is pressed if the data on that trial is particularly noisy or the decoder performs poorly in general. If the output exceeded this threshold, the feedback circle turned green and a ‘+1’ text appeared in the fixation circle, indicating that a target had been reached and a new random target would be selected. If the decoder output for the target finger was below a chosen threshold (50% probability), the feedback circle appeared grey and the target remained for the next trial. The total score (number of targets reached) was tallied and presented to participants at the end of each run. If the current target was not reached by the end of a run, that target was continued at the beginning of the next run.

### 2.5 Apparatus

Finger presses were recorded from the index, middle, ring, and little fingers of the right hand using a four-button box (Current Designs, Philadelphia, PA). The button box was affixed to a wooden board, which lay on the participant’s lap. Sandbags were placed under the right arm according to participant comfort in the first localizer session, and placed in the same position during the second and third sessions. Participants received visual instructions and visual feedback through a back-projection screen, driven by Python and PsychoPy running on a MacBook Pro.

### 2.6 fMRI acquisition

Participants were scanned in a Siemens Skyra 3T scanner with a 32-channel head coil. For all fMRI sessions, the same EPI sequence was used (TR=2 sec; 36 slices; in-plane resolution 2.3×2.3 mm; 100×100 matrix size; 2.15 mm slice thickness; 0.15 mm slice gap; 2x multiband factor). After auto-alignment to the AC-PC plane, a manual adjustment was performed to ensure full coverage of the motor cortex. The same manual adjustment parameters were applied to the subsequent localizer and neurofeedback sessions. A high-resolution T1-weighted anatomical image (MEMPRAGE; FoV 256 mm (256 x 256 matrix), 176 sagittal slices; in-plane resolution 1×1mm; 256×256 matrix size; 1 mm slice thickness; TR=2530 ms; TE=1.64/3.5/5.36/7.22) was also acquired during the first localizer session. To collect real-time fMRI data, a high-performance GPU was installed in the Measurement And Reconstruction System (MARS) that handles image reconstruction; this dedicated hardware speeds reconstruction times by more than a factor of ten. From there, the raw images are immediately sent via a 10Gb/s fiber link that runs directly from the MARS to the analysis workstation (a Dell Precision T7600n, with dual eight-core E5-2665 2.4GHz processors, 64GB 1600MHz registered ECC memory). Data is then transformed into a standard NIFTI imaging file format for preprocessing, registration, and MVPA analysis using custom Python software (Instabrain; https://github.com/LewisPeacockLab/instabrain).

### 2.7 Regions-of-Interest

Regions-of-Interest (ROIs) within sensorimotor cortex were identified using a Freesurfer (Dale et al., 1999) segmentation of the high-resolution MEMPRAGE image. Four regions were used as the basis for ROI analysis: Brodmann areas 4a, 4p, 3a, and 3b. All masks were generated simultaneously in each participant’s functional space using Freesurfer’s mri label2vol to ensure that each functional voxel was assigned to only one of the regions. These masks were then combined into the left primary motor cortex (M1: combined BA4a and BA4p) and left primary somatosensory area (S1: combined BA3a and BA3b). Five additional ROIs were then generated from combinations of these two primary ROIs: a combined ROI (M1+S1), and reduced-overlap versions (^−^and^−−^) of M1 and S1. To create these versions, the adjacent ROI (for M1: S1; for S1: M1) was expanded by one or two voxel widths (^−^: 2.3mm, ^−−^: 4.6mm, using fslmaths options: e.g. -kernel sphere 2.3 -dilM) and subtracted from the standard ROI.

### 2.8 fMRI processing

The mean of the first fMRI run of the first localizer session was used as a reference functional image (RFI). All functional volumes from both localizer sessions underwent rigid body motion correction to the RFI template using FSL’s MCFLIRT. Each functional voxel then underwent some combination of detrending and normalization (z-scoring) on a run-by-run basis. Three levels of detrending were investigated: none, real-time (using all prior data from the current run), and offline (using data from the entire run). Four types of z-scoring were investigated: none, baseline rest (using the baseline rest period at the beginning of each run), real-time (using all prior data from the current run), and offline (using data from the entire run). The baseline rest detrending condition was further subdivided to evaluate the impact of the amount of baseline data used, from 2 TRs (4 sec) up to 20 TRs (40 sec). In each localizer run there were 40 trials per finger, thus 80 total trials of data were collected per finger. Mean voxel activities were extracted for each trial using a 3-TR (6-sec) sliding window across the trial. Resulting fMRI activity patterns were then masked by ROIs and submitted to a sparse multinomial logistic regression classifier (SMLR; Krishnapuram et al. (2005)). Classifier importance maps (McDuff et al., 2009) were calculated to identify voxels that contributed significantly to the identification of each finger. For within-session analyses, a leave-one-run-out cross-validation was performed to determine decoder accuracy. For across-session analysis, the decoder was trained on one localizer session and applied to the other localizer session (and vice versa). For the neurofeedback session, real-time detrending and z-scoring were based on the full 40-sec baseline rest period. FSL’s MCFLIRT was used to realign real-time functional volumes to the RFI template for each participant. Based on offline analyses of the localizer data, we sought to maximize the quality of decoded neurofeedback by focusing on fMRI data from the combined M1+S1 ROI during the final 6-sec time window prior to feedback (TRs 4-6) on each trial.

### 2.9 Neurofeedback simulation: simulated behavior

To help calibrate design parameters and predict human performance in the neurofeedback session, we performed an offline simulation using fMRI data from the localizer sessions. The across-session fMRI decoder outputs from every trial in every localizer session were used (160 trials x 2 sessions x 6 participants = 1,920 trials, with 480 trials per finger). As in the real neurofeedback session, a pseudo-random target finger was chosen for every trial. A simple search strategy was chosen for the behavioral simulation: choosing each of the fingers sequentially, starting with the index, until the target was found (i.e., index, middle, ring, little, index-...). For each simulated pressing trial, a sample trial corresponding to the selected finger was randomly chosen (with replacement) from the full set of localizer trials to simulate the brain activity on the trial. If the decoder output for the target finger exceeded a success threshold (which could occur due to noise, leading to a false positive) then the target was considered found and a new target was chosen for the next simulated trial. Several conditions were tested: two different ROIs (M1+S1, and M1 alone), and a range of success thresholds (from 25% to 90% in 5% increments). For each combination of conditions, a total of N=1,000 participants were simulated, each performing 160 trials of simulated neurofeedback, as in the real experiment.

### 2.10 Neurofeedback simulation: human behavior

We also conducted a behavioral experiment to compare neurofeedback performance using human strategies compared to the simulated (sequential) behavioral strategy. Using the same across-session decoder outputs from the localizer sessions, a new set of human participants (N=10) attempted to find targets as in the neurofeedback experiment. The structure of the experiment was similar to the neurofeedback session except accelerated in time: participants only had to make a single press (rather than 10-sec of presses) to choose their finger, and feedback appeared immediately after the press. This accelerated timing allowed us to investigate two ROIs (using decoded neurofeedback from M1+S1 and also from M1 only) in a short period of time. We previously found no difference in trials-to-target for accelerated simulated feedback compared to the significantly slower feedback pace of real-time fMRI (Oblak et al., 2017). Furthermore, participants had no knowledge of the true target: only a one-dimensional feedback signal was provided for the target, which meant that false positives were possible, mimicking a true neurofeedback experiment in which incorrect strategies may spuriously cause positive feedback signals. We conducted 8 consecutive runs of trials for each ROI (20 trials per run, ROI order randomized across participants), similar to the real neurofeedback experiment. This experiment lasted approximated 10 minutes per participant.

### 2.11 Statistics

A separate linear mixed-effects model was created to predict decoder accuracy for each of real-time decoding, baseline sensitivity, normalization, and detrending analyses. Each model included ROI (S1 or M1) and decoding type (within or between session) as fixed effects and subject as a random effect. Tukey’s post-hoc test (α<0.05) was used to determine the differences between each condition. To compare decoding accuracy across ROIs (M1+S1, M1, S1, M1^−^, S1^−^, M1^−−^, and S1^−−^), paired t-tests (df=5) were used. To compare the correlation of decoder outputs across ROIs, the mean correlation (averaged across both localizer sessions) for each subject was Fisher Z-transformed and submitted to a paired t-test. Performance of real neurofeedback and simulated neurofeedback participants were compared using an independent groups t-test (df=13).

## 3 Results

### 3.1 Real-time decoding limitations

Using fMRI data from the two localizer sessions, we characterized the limitations of decoding individual finger presses in real-time from fMRI activity in M1 and S1 by manipulating key analysis parameters: timing, baseline duration, normalization type, and detrending type (Fig 2).

**Figure 2:**
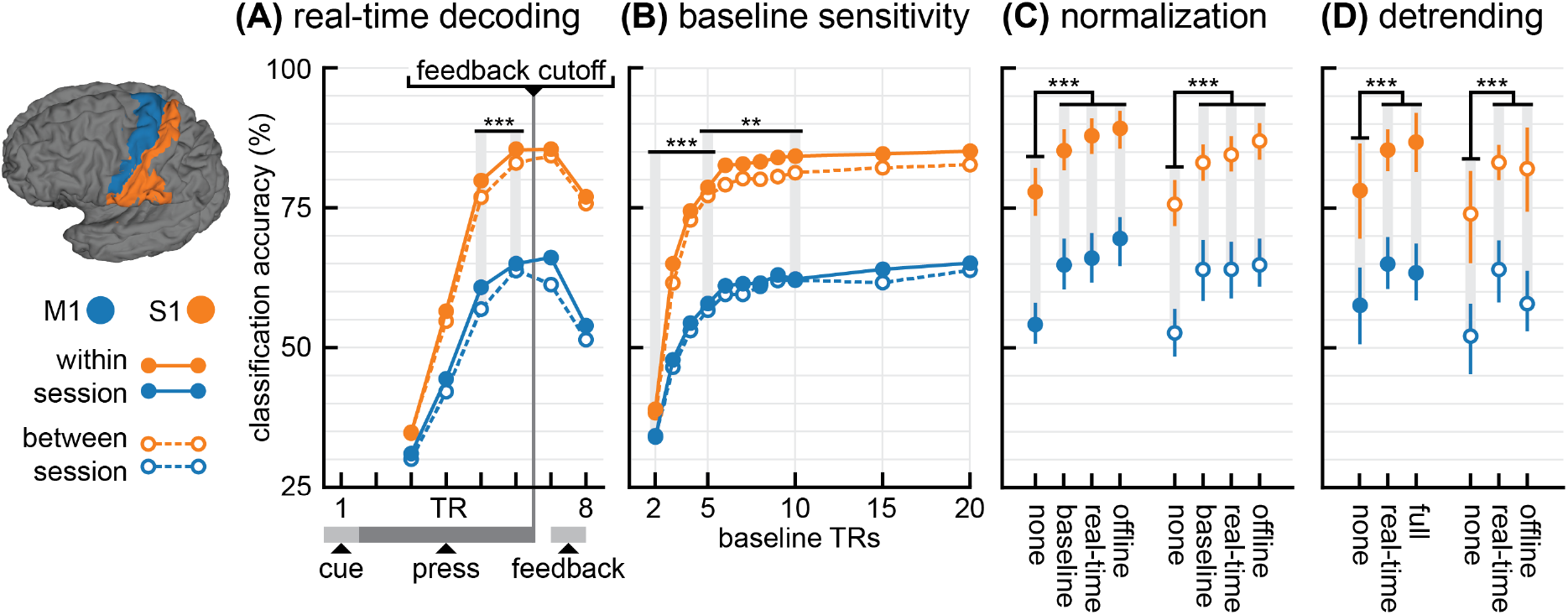
Real-time decoding limitations. **(A)** Real-time finger decoding over time (chance: 25%). The feedback cutoff (1-sec before the feedback period) is the last time at which fMRI data can be used for neurofeedback. Each displayed time point includes time-averaged fMRI data from the current time repetition (TR) and the two previous TRs. **(B)** Sensitivity of decoding to the baseline time period used for normalization (z-scoring). **(C)** Decodability for four different types of normalization: none, z-scoring based on 20 TRs of baseline data, realtime z-scoring, and z-scoring based on a full run of data, which is equivalent to offline analysis. **(D)** Decodability for three different types of detrending: none, real-time detrending, and offline detrending based on a full run of data. Error bars indicate a 95% confidence interval. M1 shown in blue and S1 shown in orange. Within session decoding shown with solid lines and closed circles and between-session decoding shown with dashed lines and open circles. Selected statistical comparisons shown. Stars indicate significant differences at p<0.01 (**) and p<0.001 (***).

#### 3.1.1 Real-time decoding over time

The timing limitations of real-time decoding were related to the intermittent feedback timing. In order to deliver feedback at the scheduled time (11 sec after the beginning of each pressing period), we could only use data gathered up to 1 sec before the feedback period. However, the timing of this feedback period could be adjusted to save time or optimize decodability. To assess the optimality of feedback timing, we analyzed decoding accuracies at TRs 5, 6, and 7, corresponding to our hypothesized optimal feedback TR (6) and the TRs immediately before and after (5 and 7). We found decoding at TR 6 to be significantly better than TR 5 (+5.6% decoding, Tukey’s HSD, p<0.001), whereas TR 7 was not significantly different than TR 6 (−0.1% decoding, p=0.996). Fig 2A illustrates these differences. This analysis also revealed a large main effect of ROI (S1 > M1: +20.1% decoding, p<0.001) and a small but reliable main effect of session (between session < within session: −2.8% decoding, p=0.004).

#### 3.1.2 Baseline sensitivity

We next investigated how sensitive the decoder was to different amounts of baseline data used for normalization (z-scoring). We analyzed four amounts of baseline data: the first 2 TRs, 5 TRs, 10 TRs, and 20 TRs of the run (Fig 2B). Increasing from using 2 to 5 TRs for baseline normalization significantly increased decodability (+31.6% decoding, p<0.001). Increasing from 5 to 10 TRs also significantly increased decodability, but with diminishing returns (+4.8% decoding, p<0.001). There was no significant difference in decodability between 10 and 20 TRs (+1.9% decoding, p=0.58).

#### 3.1.3 Normalization

Next, we analyzed how different types of normalization (z-scoring) affected decoding. We first compared baseline z-scoring (using the full 20-TR baseline period), real-time z-scoring (using all previous data from the run), and offline z-scoring (using all the data in the run, Fig 2C). Each of these was significantly better than performing no z-scoring at all (p<0.001), yielding mean decoding increases of 9.2%, 10.5%, and 12.4% decoding for baseline, real-time, and offline conditions, respectively. Within these three types of normalization, there was only a significant difference when comparing baseline to offline z-scoring (−3.3% decoding, p=0.022).

#### 3.1.4 Detrending

Finally, we investigated the effect of different types of detrending on decoding. Both real-time and offline detrending were significantly better than no detrending (p<0.001), with a mean decoding increase of 8.9% for real-time detrending and 7.2% for offline detrending (Fig 2D). There was no significant difference between real-time and offline detrending (p=0.45).

### 3.2 Decoding and information transfer across ROIs

We next investigated decoding and decoder outputs in different ROIs (Fig 3A) based on the same localizer data.

**Figure 3:**
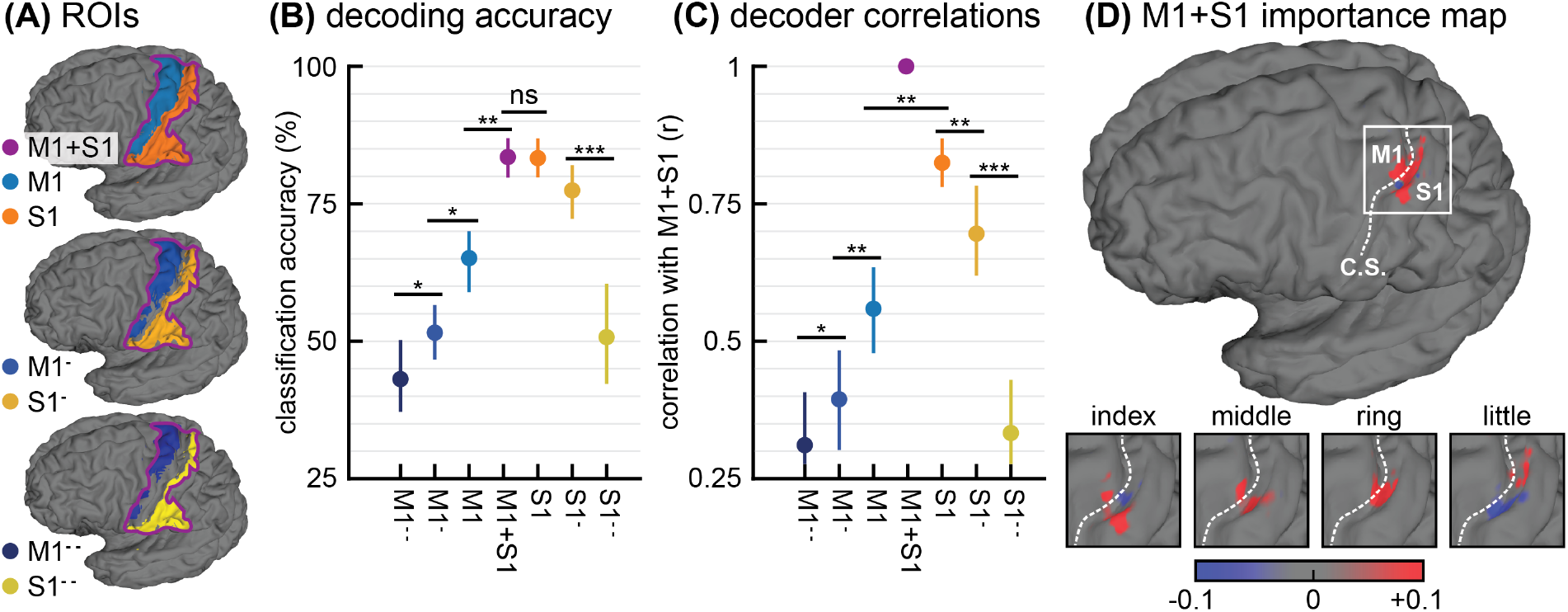
Decoding and information transfer across ROIs. **(A)** ROIs used for analysis. **(B)** Between session decoding for ROIs. (C) Correlations between decoder outputs at the same time point but in different ROIs. All correlations are made with the combined M1+S1 decoder outputs, indicating the correspondence of information between the combined M1+S1 decoder and the reduced ROIs. (D) Classifier importance maps for the combined M1+S1 decoder of a sample participant, in arbitrary units. C.S. indicates the central sulcus dividing M1 and S1. Error bars indicate a 95% confidence interval. Selected statistical comparisons shown. Stars indicate significant differences at p<0.05 (*), p<0.01 (**), and p<0.001 (***). ‘NS’ indicates no significant differences.

#### 3.2.1 Decoding accuracy

Across-session decoding accuracy was highest in S1 (83.4%) and in the combined M1+S1 region (83.3%), with no significant difference between the two (p=0.85). M1+S1 had significantly better decoding than M1 (+18.2% decoding; p=0.002). Moving to the reduced M1 from M1 was significantly worse (−13.8%, p=0.019), and also to M1^−−^ from M1^−^ (−8.2%, p=0.027). S1^−^ was not significantly worse than S1 (−5.94%, p=0.073), but S1^−−^ was significantly worse than S1^−^ (−26.7%, p<0.001). Fig 3B illustrates differences in decoding accuracy.

#### 3.2.2 Decoder correlations

The decoder outputs from S1 were more strongly correlated with the combined M1+S1 region than were the decoder outputs from M1 correlated to this region (p=0.005). Moving anteriorly, the M1^−^ decoder was more weakly correlated than M1 (p=0.003), and M1^−−^ was more weakly correlated than M1^−^ (p=0.016). Similarly, moving posteriorly, the S1 decoder was more weakly correlated than S1 (p=0.003), and S1^−−^ more weakly correlated than S1^−^ (p<0.001). See Fig 3C for correlation results.

#### 3.2.3 Importance maps

Classifier importance maps (McDuff et al., 2009) show that important voxels lie on both sides of the central sulcus, with more on the posterior (S1) side (Fig 3D). Furthermore, they tend to be near to the central sulcus.

### 3.3 Finger finding experiment

#### 3.3.1 Predicted performance

We first predicted performance based on a variable success threshold for finding targets in both the combined M1+S1 region and in the M1 region alone (Fig 4A). Increasing the success threshold caused an exponential increase in the number of trials required to find a target, with diminishing returns on predicted target accuracy. Based on these predictions, a threshold of 50% was selected for subsequent experiments with human participants to maximize accuracy while minimizing the number of trials to target.

**Figure 4:**
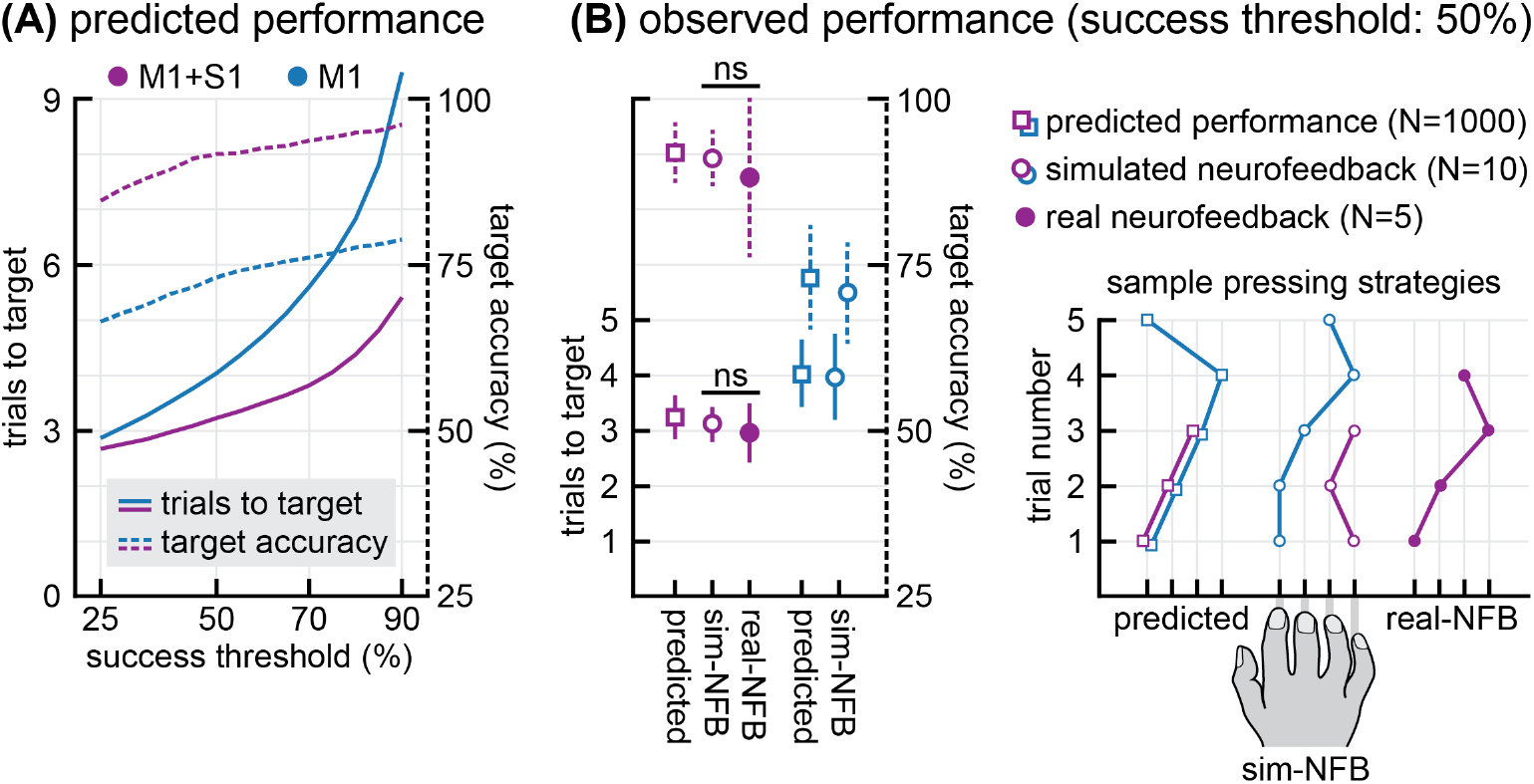
Finger finding neurofeedback experiment performance. **(A)** Predicted performance based on between-session decoder outputs from the localizer sessions. The success threshold for finding targets was varied between 25% and 90%. Target accuracy indicates the proportion of targets in which the finger pressed when the decoder output exceeded the success threshold matched the target finger. **(B)** Observed performance for both real neurofeedback and simulated neurofeedback participants using a success threshold of 50%. Trial-by-trial sample pressing strategies are shown for each condition: predicted, simulated, and real neurofeedback. Each tick represents a single finger of the right hand as illustrated. Each line represents one example target search, with the line ending when the decoder output exceeded the target threshold. Due to noise, occasionally the same finger is repeated. Error bars indicate standard deviation. M1 shown in blue and M1+S1 shown in purple. ‘NS’ indicates no significant differences.

#### 3.3.2 Simulated neurofeedback results

We next investigated whether human participants’ performance matched our model predictions for the selected success threshold (Fig 4B). In M1+S1, we expected 3.23+/-0.29 trials to target (mean+/-s.d.) and 91.7+/-3.7% target accuracy. In our simulated neurofeedback experiment, we recorded 3.12+/-0.12 trials to target with 91.9+/-3.4% accuracy using data from this region. In the M1 region, our predictions were similarly accurate: the prediction was 4.04+/-0.49 trials with 73.1+/-7.1% accuracy, and the simulated neurofeedback result recorded was 3.97+/-0.67 trials with 70.1+/-6.8% accuracy.

#### 3.3.3 Real neurofeedback results

We then compared real neurofeedback results to the simulated results in the M1+S1 region that was selected for the neurofeedback experiment (Fig 4B). In the scanner, participants required 2.97+/-0.43 trials to find each target, with 87.7+/-11.7% accuracy. There was no significant difference between this performance and the simulated neurofeedback participants’ performance (trials-to-target: t_(13)_=-0.97, p=0.35; accuracy: t_(13)_=-0.78, p=0.45).

Finally, we investigated how decoding accuracy influenced performance on our task (Fig 5). We selected the 50% success threshold and M1+S1 region in order to qualitatively compare performances between the three conditions (predicted, simulated neurofeedback, and real neurofeedback). In the predicted dataset, trials-to-target was negatively correlated with decoding accuracy (slope of best-fit line=-4.36, r=-0.41, p<0.001 Fig 5A) and target accuracy was positively correlated with decoding accuracy (slope=0.66, r=0.48, p<0.001, Fig 5B), as expected. The slope for the trials-to-target was negative for simulated neurofeedback (slope=-0.86) and real neurofeedback (slope=-3.19); the slope for decoding accuracy was positive for simulated neurofeedback (slope=0.70) and real neurofeedback (slope=0.93).

## 4 Discussion

This study presents a systematic investigation of optimal parameter selection for the design of real-time neurofeedback experiments that rely on multi-voxel pattern analysis of neuroimaging data. We collected fMRI data of participants performing individual finger presses and trained classifiers to discriminate brain activity patterns for each finger in sensorimotor cortex. These classifiers were intended to perform real-time decoding of finger presses in a subsequent neurofeedback session. Offline analyses with real-time processing constraints revealed optimal choices for the analysis of these data and the timing of the neurofeedback experiment. Simulated results of neurofeedback performance were confirmed by participants receiving real neurofeedback inside the scanner. This study demonstrates how the offline simulation of neurofeedback performance using real fMRI data can be used to optimize human performance on real-time neurofeedback experiments.

**Figure 5:**
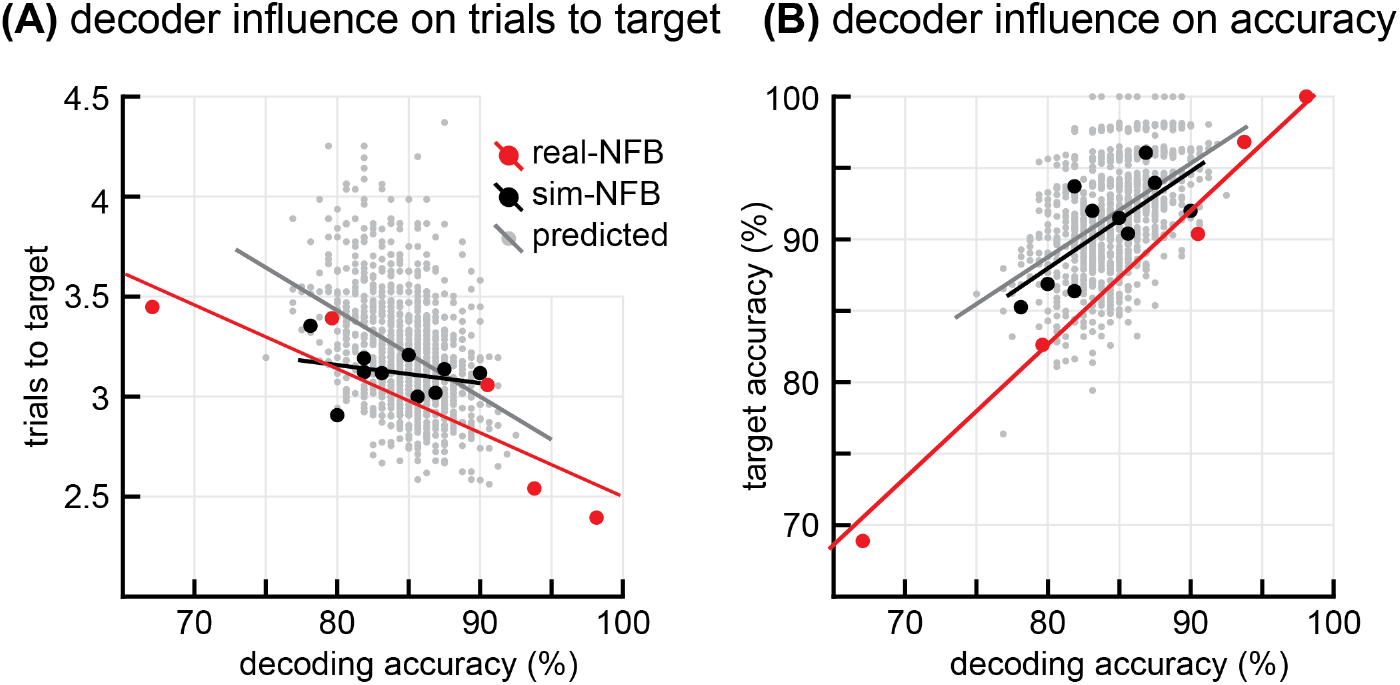
Influence of decoding accuracy on finger finding performance. **(A)** Correspondence between observed decoding accuracy during the finger finding session and the mean number of trials required to find each target. **(B)** Correspondence between observed decoding accuracy during the finger finding session and target accuracy (proportion of trials in which the decoder output exceeded the success threshold and the pressed finger was also the target finger). Solid lines indicated the best-fit line for each condition.

The simulation of neurofeedback performance is a novel approach to the field. While reliance on simulations is common in other neuroscience subdomains such as visual neuroscience (Tong, 2003), the only other instance of simulation in fMRI neurofeedback is our previous work mentioned earlier (Oblak et al., 2017). Whereas in the aforementioned study, we simulated brain activity based on known parameters of visual cortex activity, here we present recorded brain activity in sensorimotor cortex to both to human participants and to simulated participants with predetermined neurofeedback strategies. A key element to this simulation of neurofeedback is an explicit strategy, i.e. finger pressing, because it can be directly measured and validated both inside and outside of the scanner. Our simulation was validated for explicit strategies through similar outcomes in real and simulated neurofeedback (Fig 4B). However, it remains to be answered to what degree the explicit strategies used in this paradigm can represent mental strategies typically used in neurofeedback. Future work should incorporate more advanced learning models, including implicit strategies, into this simulation paradigm. We anticipate such simulation of neurofeedback performance to be a valuable tool to improve the efficiency and robustness of experiments.

Our work shows the benefit of using localizer data to predict neurofeedback performance in two ways. First, we were able to design a parameter of our experiment, namely the feedback success threshold of 50%, to optimize predicted neurofeedback performance as a tradeoff between finding successful strategies and finding accurate strategies (Fig 4A). These predictions translated to both real neurofeedback participants and human participants interacting with a simulated neurofeedback signal outside of the fMRI scanner (Fig 4B). Although our task and model of learning was simple, this same simulation strategy could be used with a more complex learning model (Oblak et al., 2017; Watanabe et al., 2017) to determine the required duration of an experiment, or to design tuning curves in an adaptive neurofeedback experiment (deBettencourt et al., 2015). Second, given our fully designed experiment, we were able to predict the neurofeedback performance of individuals (Fig 5). This type of prediction could be used as an exclusionary criterion for patients in neurofeedback treatment or to predict the required duration of treatment; it could also be used to alter neurofeedback parameters for individuals based on their own localizer data.

Our neurofeedback predictions show how a given decoder accuracy translates to neurofeedback performance. In all cases, neurofeedback performance increased with increasing decoder accuracy. Therefore, our secondary goal was to explore which typical parameters of decoded neurofeedback experiment had the largest effect on decoding accuracy. Two standard fMRI preprocessing steps, normalization and detrending, were found to have a large effect on real-time decoding. Real-time z-scoring increased decoding accuracy by 10.5% (Fig 2C) and real-time detrending increased decoding accuracy by 8.9% (Fig 2D) compared to no preprocessing at all. Critically, these results did not suffer relative to offline decoding, indicating that the real-time preprocessing constraints were not a performance bottleneck.

We next found that we could decode from an earlier timing window than standard decoded neurofeedback experiments without a significant reduction in decoder accuracy. The standard timing window for decoded fMRI neurofeedback is a 6-sec stimulus period followed by a 6 sec decoding period (Shibata et al., 2011), accounting for the hemodynamic delay. We increased the length of the stimulus period (in this case, finger pressing) to 10 sec, yet decoded 2 sec earlier than Shibata et al. (2011) (Fig 2A). For behaviors and ROIs other than those detailed in this work, we recommend designing a localizer with varying stimulus periods and rest periods to determine the best tradeoff between decodability and feedback timing for that experiment

The standard normalization method for decoded neurofeedback is z-scoring using a baseline resting period of 20 sec at the beginning of each fMRI run (Shibata et al., 2011). Our results support this choice, as there was no significant increase in subsequent decodability when we doubled the length of the baseline period to 40 sec. However, we also show that real-time z-scoring can be used to achieve similar decoding performance (Fig 2B), calling into question the necessity of this baseline period. The success of real-time z-scoring may have been due to our study having a strict set of strategies (one of four finger presses) for participants. In less constrained situations, results may depend on the variability of participants’ strategies (and neural activity) during neurofeedback trials. However, it would remain valid to normalize based on a baseline rest period each run.

We observed differences in decoder performance dependent on the specificity and location of the ROI. For instance, we found that by reducing the size of our ROIs by only one voxel width, decoding accuracy was reduced by 14% in M1 and by 6% in S1 (Fig 3B). These results are not surprising given the limited spatial resolution of fMRI and high intersubject variability. This evidence suggests that predetermined segregation of ROIs should be handled carefully. If a neurofeedback experiment targets a specific behavior without a well-defined anatomical hypothesis, then we should err on the side of inclusion and allow the decoder to automatically select relevant voxels in the brain (Shibata et al., 2016). However, if there is a strict hypothesis based on a neural mechanism in a specific ROI, then it is reasonable to restrict the voxels at the expense of decodability (Shibata et al., 2011). In our case, we chose a broad M1+S1 ROI because we were not attempting to segregate a specific neural mechanism, such as separating motor output from tactile sensation. If we had a strict motor or sensory hypothesis, isolating the M1 or S1 ROI may be necessary. However, if such segregation was necessary, and subsequently reduced decoder performance, the procedures illustrated here could be used to predict neurofeedback performance, such as the number of trials required to induce neurofeedback learning. Fig 5 shows how decoder accuracy is predictive of neurofeedback performance. Thus, one of the contributions of the of this work is the suggestion that a targeted decoder accuracy should be predetermined together with the goals of the study prior to commencing neurofeedback training.

While here we show the power of simulation of decoded neurofeedback parameters, there are limitations to what we can conclude. For example, the number of neurofeedback participants was low (N=5), especially compared to the simulated neurofeedback participants (N=10). As expected, we did not find a difference in performance between the two groups, validating our model. It is possible, however, that a difference may arise with a larger sample of neurofeedback participants. If differences were found, the model could be improved to address more subtle facets of the experiment. For example, although participants were not instructed to self-modulate their brain activity, their performance could nonetheless be affected by neurofeedback-induced changes in the brain. If real neurofeedback performance were to diverge from simulated performance, this is one likely factor that should be accounted for in the model. Perhaps more importantly, we focused only on M1 and S1, which not only could affect our model, but also generalizability of the model to other brain regions. Another mitigating factor is the composition of variability in decoder output: we cannot conclude whether the variability in decoder output is due to measurement noise, spontaneous neural activity, or variability in motor behavior. Assessing the source of variability in decoder output is a key component of modeling decoded neurofeedback that future work must address.

### 4.1 Conclusions

In this work we show that decoded neurofeedback performance is highly correlated to decoder accuracy, and we systematically determine the parameter settings needed to optimize that decoder accuracy. We modeled neurofeedback performance using simulations based on real brain data, compared this with human performance with the same brain data, and finally compared it with real neurofeedback performance. We observed similar performance in all cases, validating the accuracy of the simulations. We found a quantitative representation of the high level of dependence of neurofeedback performance on success threshold and decoder accuracy. These results will help improve the robustness of decoded neurofeedback experiments, both by identifying proper preprocessing parameters and by identifying likely ‘non-responder’ participants prior to training. The simulation paradigm validated here can be used in future research to pre-emptively and efficiently sweep the parameter space to optimize the design of decoded eurofeedback experiments.

